# Contrasting patterns of coding and flanking region evolution in mammalian keratin associated protein-1 genes

**DOI:** 10.1101/282418

**Authors:** Huitong Zhou, Tina Visnovska, Hua Gong, Sebastian Schmeier, Jon Hickford, Austen R.D. Ganley

## Abstract

DNA repeats are common in eukaryotic genomes, and recombination between copies can occur. This recombination can result in concerted evolution, where within-genome repeats are more similar to each other than to orthologous repeats in related species. We investigated the tandemly-repeated keratin-associated protein (KAP) gene family, *KRTAP1*, which encodes proteins that are important components of hair and wool in mammals. Comparison of *KRTAP1* gene repeats across the mammalian phylogeny shows strongly contrasting evolutionary patterns between the coding regions, that have a concerted evolution pattern, and the flanking regions, that have a normal, radiating pattern of evolution. This dichotomy transitions abruptly at the start and stop codons, and is not the result of purifying selection, codon adaptation, or reverse transcription of *KRTAP1-n* mRNA. Instead, our results suggest that short-tract gene conversion events coupled with selection for these events in the coding region drives the contrasting *KRTAP1* repeat evolutionary patterns. Our work shows the power that repeat recombination has to complement selection and shape the evolution of repetitive genes, and this interplay may be a more common mechanism than currently appreciated for achieving adaptive outcomes in eukaryotic multi-gene families. Thus, our work argues for greater emphasis on exploring the evolution of these families.

## Introduction

Repetitive DNA is widespread in most eukaryote genomes (Britten and Kohne 1968; Richard, et al. 2008; Lopez-Flores and Garrido-Ramos 2012). There are two basic repeat DNA types: tandem repeats that are typically arranged in head-to-tail arrays; and dispersed repeats, and these can occur in either coding or non-coding DNA. Repeats are thought to arise from recombination-based duplication/amplification events (Stephan 1989). Sequence identity between duplicates will then decay through the diversifying force of mutation, unless counteracting processes operate (Brown, et al. 1972; Dover 1982). The balance between duplication, diversification, selection, and counteracting forces thus dictate the evolutionary dynamics of repeats. Two main paradigms have been proposed to account for the long-term maintenance of repeat identity: concerted evolution and birth-and-death evolution. Concerted evolution describes a pattern of evolution where the repeats within a genome show greater sequence identity to each other than to orthologous repeats in related genomes (Elder and Turner 1995). The pattern of concerted evolution is proposed to result from recombination-based processes, such as gene conversion and unequal cross-over events, that replace the DNA sequence from one repeat with that from another repeat (Liao 1999). In so doing, these recombination processes maintain sequence identity between repeat copies in the face of mutation, and thus homogenize the repeats (Dover 1982). ‘Birth-and-death’ evolution involves purifying selection maintaining sequence identity between repeats that are generated by occasional duplication events (i.e. birth), as well as death, which results from repeat loss or pseudogenization (Nei, et al. 1997; Nei, et al. 2000). While there has been debate as to which of these processes best describes the evolutionary dynamics of repetitive DNA (Nei and Rooney 2005; Rooney and Ward 2005; Eirin-Lopez, et al. 2012), a basic characterization of the evolutionary dynamics of most repeat families is lacking.

The keratin-associated proteins (KAPs) are a diverse group of proteins, and are rich in either sulphur, or glycine and tyrosine. They are important structural components of hair and wool fibres, and form a matrix that cross-links the keratin intermediate filaments. The genes encoding the KAPs are called *KRTAP*s (Gong, et al. 2012), and can be classified into 27 families, with each family comprising 1-12 members that are usually tandemly arranged (Rogers and Schweizer 2005; Rogers, et al. 2006; Gong, et al. 2016). The *KRTAP*s are single exon (intron-less) genes, with small coding sequences (less than 1 kb) (Rogers and Schweizer 2005), and they have low numbers of pseudogenes. For example, in humans the pseudogene:gene *KRTAP* ratio is approximately 1:5 (Gong, et al. 2016), while across all human genes the ratio is close to 1:1 (Torrents, et al. 2003; Stein 2004). In addition, the *KRTAP*s show high levels of population variation, with all known *KRTAP* genes being polymorphic in sheep (Gong, Zhou, McKenzie, et al. 2010; Gong, et al. 2016; Zhou, et al. 2016), where they are well studied because of their roles in determining wool phenotypes (Zhou, et al. 2015; Li, Zhou, Gong, Zhao, Hu, et al. 2017; Li, Zhou, Gong, Zhao, Wang, Liu, et al. 2017; Li, Zhou, Gong, Zhao, Wang, Luo, et al. 2017; Tao, Zhou, Gong, et al. 2017; Tao, Zhou, Yang, et al. 2017). Despite this variation, it has been reported that at least some *KRTAP* genes show a pattern of concerted evolution between the paraglogous gene copies (Rogers, et al. 1994; Wu, et al. 2008; Khan, et al. 2014).

The KAP1 proteins form the best characterised KAP family, and they show a high degree of sequence heterogeneity compared to other KAP families. These KAP1 proteins appear to be restricted in expression to the middle to upper cortex region of the hair and wool follicle, and are absent in the cuticle (Powell and Rogers 1997; Shimomura, et al. 2002). Their precise role in hair and wool function, has yet to be determined. The genes encoding the KAP1 proteins (*KRTAP1- n*) have been characterized in a number of mammalian species, where they are usually arranged as four tandem copies (**Figure 1**) (Khan, et al. 2014). The coding regions of the *KRTAP1-n* genes vary in length within species, predominantly as a consequence of variation in the number of imperfect tandem decapeptide repeat units (Gong, et al. 2016) (**Figure 1**).

**Figure 1.**
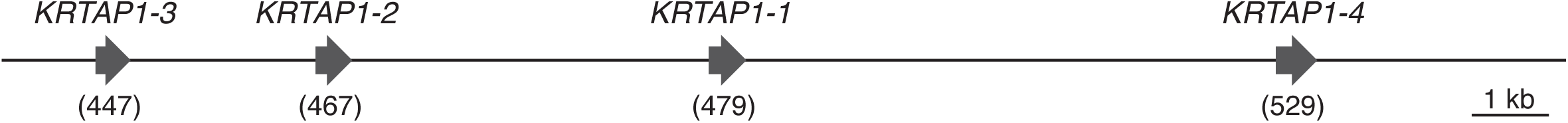
Tandem repeat organization of the keratin associated protein-1 (*KRTAP1*) genes. The organization of mammalian *KRTAP1* genes is illustrated by the arrangement found in sheep. The four *KRTAP1-n* paralogs are represented by arrows that indicate the direction of transcription. Diagram is drawn to scale, with *KRTAP1-n* lengths bracketed below the genes. These repeats are numbered *KRTAP1-1, 3, 4*, and *5* in human.

Here we analyse the *KRTAP1* genes from a number of mammalian species, including four species for which the *KRTAP1-n* loci have not been described. Together with the existing *KRTAP1-n* sequences, we reveal that the *KRTAP1-n* coding regions display a pattern of concerted evolution. In stark contrast to the coding region though, we find that the repeat flanking regions display no evidence of concerted evolution, and instead appear to be evolving by normal vertical or radiating evolution. Surprisingly, we find that this pattern of coding region restricted concerted evolution is not the result of purifying selection, nor does it result from codon adaptation or reverse transcription/reintegration of *KRTAP1-n* mRNA sequences. Instead, the results are best explained by a combination of on-going short-tract gene conversion events between the *KRTAP1-n* copies, and negative selection. We argue that these gene conversion events act as an unusual mechanism of purifying selection to prevent excessive intra-genomic divergence between the four gene copies, while also allowing inter-species diversity. This unusual mode of evolution may apply to other multicopy genes that encode products subject to diversifying selection.

## Materials and Methods

### Sequence Resources and Gene Identification

All genome sequences were sourced from the NCBI GenBank. Previously identified *KRTAP1-n* sequences (Itenge-Mweza, et al. 2007; Wu, et al. 2008; Gong, Zhou and Hickford 2010; Gong, et al. 2011) were used to search the genomes of cattle, horses, rabbits and African elephants using BLAST with default parameters, and the genes retrieved were identified by sequence identity within both the coding and flanking regions (**Table S1**).

### Sequence Alignments

*KRTAP1* nucleotide sequences (**Table S1)** for all four paralogs from the ten species (sheep, cattle, dog, elephant, horse, human, macaque, mouse, rat and rabbit) were separated into 5’ flanking regions, coding sequences, and 3’ flanking regions. The multiple sequence alignment tool *mafft* (v7.123b) (Katoh and Standley 2013) was used to separately align the 5’ and 3’ flanking regions as nucleotide sequences, using the arguments ‘--nuc --localpair --maxiterate 1000’. To align the coding sequences at the predicted amino acid level, *mafft* with the arguments ‘--amino --localpair --maxiterate 1000’ was run.

The coding sequence alignment was subsequently reverse translated using *revTrans* (v1.4) (Wernersson and Pedersen 2003) with two input files: the sequences of all the coding regions, and the amino acid sequence alignments. The sequences in the two files were paired by name using the ‘-match name’ parameter, and default values were used for all other parameters. A number of regions align poorly and have many indels, therefore we used the longest continuous coding sequence block (198 nucleotides; covers on average around 40% of the coding region) where none of the 40 sequences had indels. For the flanking region alignments, we used *Gblocks* (v0.91b) (Talavera and Castresana 2007) to select blocks that cover approximately 40% of the flanking regions having the best alignment. We also used *Gblocks* with less stringent criteria to create multiple sequence alignments of the coding and flanking regions that included more poorly aligning regions.

### Phylogenetic Trees

*PhyML* (v3.1) (Guindon, et al. 2010) was used to construct phylogenies based on the coding and flanking region sequences. The number of resampled bootstrap data sets was set to 1000 (parameter ‘-b 1000’), and the additional arguments ‘-q -s BEST -o tlr’ were employed. The Bioconductor package *ggtree* (v1.9.4) (Yu, et al. 2017) was used to plot the phylogenies.

### Codon Adaptation Index

The CAIcal server (http://genomes.urv.es/CAIcal(Puigbo, et al. 2008) was used to calculate CAI values for the *KRTAP1*s, as well as expected CAI values from permutated sequences using default parameters and published codon usage data (Nakamura, et al. 2000).

### Motifs in the Coding Sequences

We used MEME motif finder (v4.12.0) (Bailey, et al. 2006) to explore repetitive elements in the coding sequences. The repetitive structure of the coding regions reported in the Results was obtained with parameters ‘-dna -oc. -nostatus -time 18000 - maxsize 60000 -mod anr -nmotifs 6 -minw 6 -maxw 30 -minsites 20 -maxsites 600 –revcomp’ and all the other parameters set to the default values.

### *KRTAP1-n* Polymorphism in Sheep

Intra-specific variation was assessed using three sequences for *KRTAP1-1* (Itenge-Mweza, et al. 2007), eleven sequences for *KRTAP1-2* (Gong, et al. 2011; Gong, et al. 2015), nine sequences for *KRTAP1-3* (Itenge-Mweza, et al. 2007), and nine sequences for *KRTAP1-4* (Gong, Zhou and Hickford 2010). These were aligned using DNAMAN (v5.2.10; Lynnon BioSoft, Canada) with default parameters, and polymorphic sites were identified manually.

### Data Availability

Sequence data are available at GenBank and the accession numbers and positions are listed in the **Materials and Methods** (sheep polymorphism data) and Table S1 (*KRTAP1* sequences).

## Results

### Mammalian *KRTAP1-n* repeats show a concerted evolution pattern in the coding but not the flanking regions

To better understand the genetic architecture of the mammalian *KRTAP1* cluster, we selected the *KRTAP1* genomic region from key members of the mammalian phylogeny for analysis. The Basic Local Alignment Search Tool (BLAST) was used to search GenBank with known *KRTAP1-n* sequences to identify and retrieve the *KRTAP1* clusters from the genomes of four species (cattle, horses, rabbits and African elephants) for whom *KRTAP1-n* sequence information has not been reported (**Figure S1**). We then combined these with previously-identified *KRTAP1- n* sequences from other mammalian species to obtain sampling across the mammalian phylogeny (**Figure 2**).

**Figure 2.**
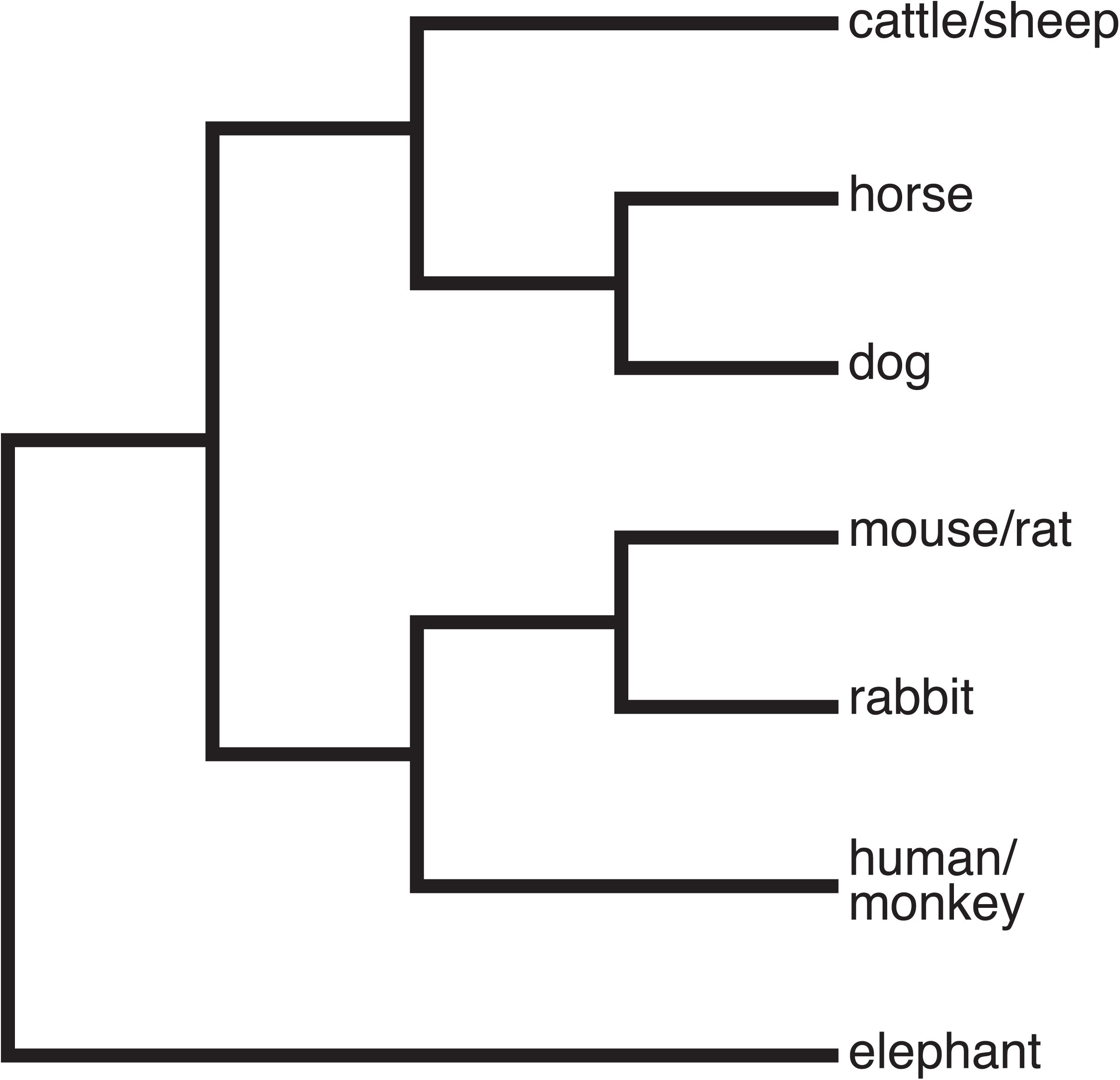
Mammalian *KRTAP1-n* gene phylogenetic relationships. Representative phylogenetic tree illustrating the relationships between the *KRTAP1-n* genes in the species used in this study. Branch lengths are not to scale. The phylogeny is adapted from that presented in McCormack et al. (2012).

Previously, the *KRTAP1* genes of sheep were shown to contain a variable number of occurrences of a QTSCCQPXXX decapeptide tandem repeat in the N-terminal region of the protein (Rogers, et al. 1994; Gong, et al. 2011; Gong, et al. 2016). We used a motif finding tool (MEME; (Bailey, et al. 2006) to search for repetitive motifs in the coding regions of all the mammalian *KRTAP1-n* sequences. This revealed that the decapeptide repeat is present at the N-terminus in all mammalian *KRTAP1-n* genes we obtained (**Figure S2)**, albeit with less amino acid conservation than that observed in sheep. MEME also identified nucleotide level tandem copies of this repeat at the C-terminus of the protein. Furthermore, both the N- and C-terminal repeats vary in copy number, within and between genomes. This copy number variation is responsible for much of the length variation between *KRTAP1-n* sequences.

To determine the genetic relationships between of the mammalian *KRTAP1-n* genes, we generated a *KRTAP1* phylogenetic tree from an alignment of our mammalian *KRTAP1-n* coding region sequences. This revealed that, in most cases, the *KRTAP1* genes are more related to each other within a species than to their orthologs in other species, thus exhibiting a concerted evolution pattern. This manifests as clades that group by species, rather than by repeat, in the phylogenetic tree (**Figure 3**). This concerted evolution pattern breaks down between the most closely-related species pairs (cattle/sheep, rat/mouse, human/macaque), presumably because the signal is confounded by these species having more recent shared ancestry. Nevertheless, for most species there is a clear pattern of concerted evolution.

**Figure 3.**
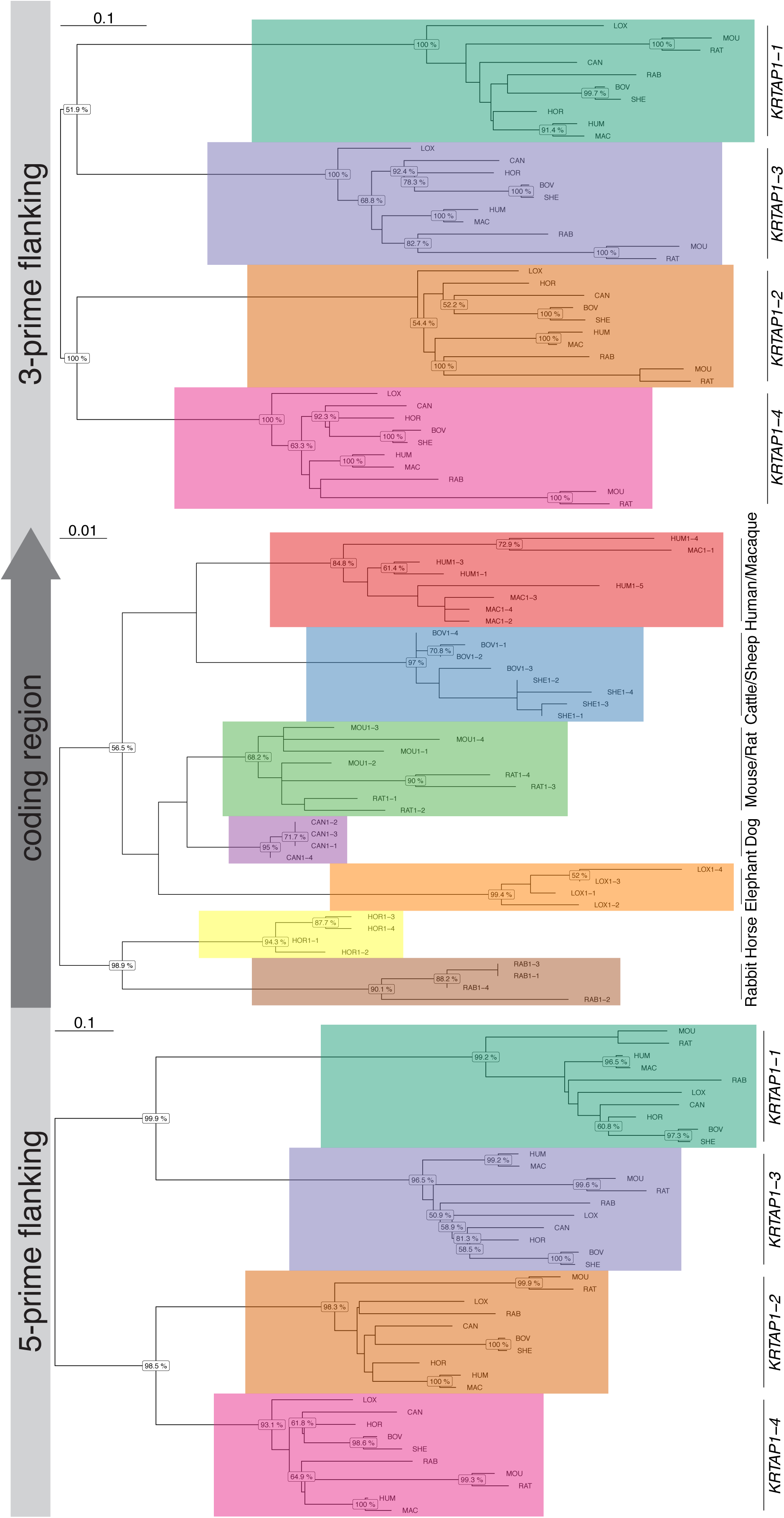
Phylogenetic trees of *KRTAP1-n* coding and flanking region sequences. Phylogenetic trees were constructed for the mammalian *KRTAP1-n* 5’ flanking region, coding region, and 3’ flanking region using PhyML. The species are indicated by three-letter abbreviations. The number following this for the coding regions indicates the *KRTAP1-n* gene name. The major clades within the trees are indicated by coloured boxes. The 5’ and 3’ flanking region phylogenies group by repeat number, while the coding region phylogeny tends to group by species. Numbers on nodes indicate bootstrap supports over 50%, and substitution rates are indicated at the top left. Human *KRTAP1-n* gene names have been altered for consistency with other species.

For concertedly evolving tandem repeat sequences such as the ribosomal RNA gene repeats, homogenization occurs for the complete repeat unit, including the non-coding regions (Ganley and Kobayashi 2007). To test whether the *KRTAP1* clusters display a ‘whole-unit’ pattern of concerted evolution, we generated *KRTAP1* phylogenetic trees from multiple alignments of the 5’ and 3’ flanking sequences of the mammalian *KRTAP1* genes. Surprisingly, the phylogenies derived from these flanking sequences did not show any pattern of concerted evolution, and in contrast to the coding region phylogeny, the clades in these phylogenetic trees were group by *KRTAP1* repeat number, not by species (**Figure 3**). We note that bootstrap support is not strong for all the clades in these phylogenetic trees, but the contrast between the coding region concerted versus flanking region radiating evolutionary patterns is unmistakable. Furthermore, the topology within many of the *KRTAP1* flanking region clades is consistent with the reported mammalian phylogeny (refer to **Figures 2** and **3**). These phylogenies were generated from multiple sequence alignments that encompass the regions that align well, but phylogenies derived from sequence alignments that include poorly aligned regions give qualitatively similar results (**Figure S3**). Overall, in stark contrast to the coding region, the flanking regions show a phylogenetic pattern expected for normal radiating evolution, and exhibit no evidence of concerted evolution.

### What is responsible for the different evolutionary patterns of the *KRTAP1* coding and flanking regions?

The difference in evolutionary pattern between the coding and flanking regions is striking, hence we sought to identify the mechanism(s) responsible.

### Purifying selection

Previous studies have shown that multi-gene loci undergoing birth-and-death evolution can show high levels of identity within the coding region due to strong purifying selection (Nei, et al. 2000; Piontkivska, et al. 2002). It is possible that purifying selection maintains sequence identity between *KRTAP1-n* copies within a species, whilst diversifying selection results in differences between species. If so, we would predict that while the non-synonymous sites would show a concerted evolution pattern, the synonymous sites would instead show a normal radiating pattern of evolution (resembling the flanking regions).

To investigate this, we looked at the pattern of evolution of the synonymous sites in the coding sequences compared to the non-synonymous sites. The number of KAP1 amino acid changes present within and between species makes it difficult to consistently call sites as synonymous or non-synonymous, so third codon positions were used as a proxy for synonymous sites, and first and second codon positions were used as a proxy for non-synonymous sites. We generated phylogenetic trees from multiple sequence alignments of the first-second (which we refer to as “non-synonymous”), and third (which we refer to as “synonymous”) codon sites of the *KRTAP1- n* coding regions to test for different evolutionary patterns. Surprisingly, while the non-synonymous sites displayed a pattern of concerted evolution as was expected (**Figure 4A**), the synonymous sites also revealed the same pattern of concerted evolution (**Figure 4B**). The concerted evolution pattern for the synonymous sites seems to be stronger than that of the non-synonymous sites, as they separate sheep and cattle into separate clades, and also resolve dog, elephant, and rat/mouse into separate clades (**Figure 4**).

**Figure 4.**
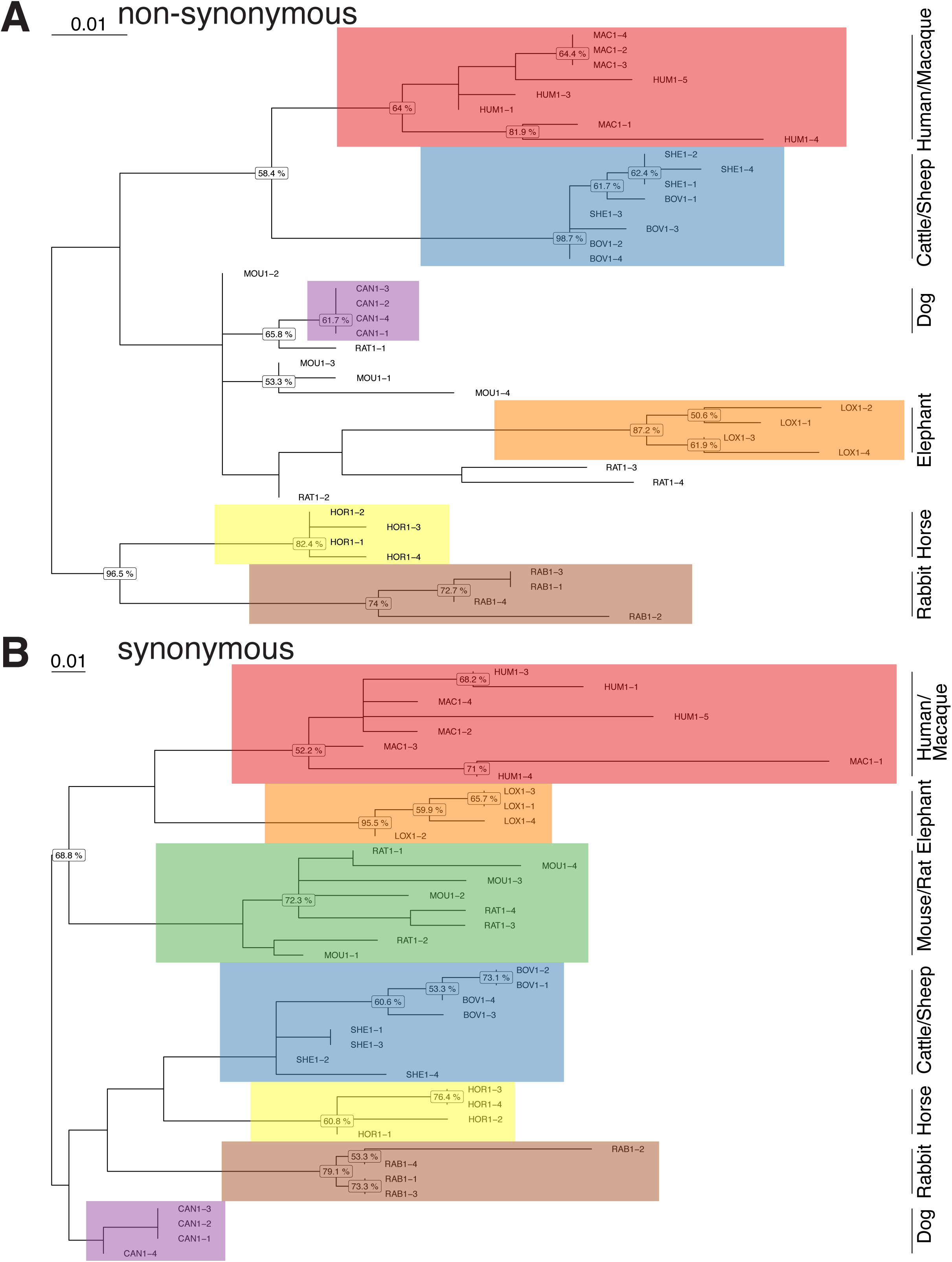
The *KRTAP1-n* concerted evolution pattern is not explained by purifying selection. Phylogenetic trees were constructed for the 1st and 2nd codon positions (“non-synonymous”; **A**), and the 3rd codon position (“synonymous”; **B**), as per **Figure 3**. The major clades in both phylogenies tend to group by species, with this concerted evolution pattern being stronger for the synonymous phylogeny. Numbers on nodes indicate bootstrap supports with values over 50%, and substitution rates are indicated at the top left.

### Codon adaptation

We considered whether this pattern of concerted evolution amongst the synonymous sites might result from codon adaptation (Lin, et al. 2006), as a result of synonymous mutations being selected to follow changes in the favoured codons between species. The *KRTAP1-n* genes display strong evidence for codon adaptation (the degree to which the favoured codons for that species are used in a gene). For example, the human *KRTAP1-n* genes collectively show a codon adaptation index (CAI) of 0.91 (out of a maximum of 1), higher than the CAI of randomly permutated human *KRTAP1* sequences (CAI=0.78). Using the *KRTAP1* coding sequence alignment used for the phylogenies presented in **Figure 3**, we identified nine synonymous differences between human and mouse that exhibit a concerted evolution pattern (similarity within species versus difference between species). If codon adaptation can explain this pattern, these synonymous mutations should change in a manner consistent with a change in codon usage preference for that amino acid. Five of these mutations show the pattern expected, given the change in codon usage between human and mouse (synonymous change creates the more favoured codon in the species it is found in). However, four of these mutations show the opposite pattern, and most of the codon usage preference changes between human and mouse are small (**Table S2**). These results provide no evidence for adaptation to different codon usage preferences driving the pattern of *KRTAP1* concerted evolution.

### Reverse transcription of *KRTAP1* mRNA

Another potential explanation for the incongruence in evolutionary pattern between the *KRTAP1* coding and flanking regions is reverse transcription of *KRTAP1-n* mRNAs, followed by homologous recombination-mediated replacement of a genomic *KRTAP1-n* with the reverse transcribed copy (Coulombe-Huntington and Majewski 2007). This is feasible given that *KRTAP1-n* are single-exon genes. If reverse transcription events occur, the 5’ and particularly 3’ flanking regions should show a concerted evolution pattern that is similar to the coding region. Inspection of the 5’ and 3’ flanking regions revealed that sequence similarity between *KRTAP1-n* sequences within a genome tends to decay immediately upstream of the ATG codon and downstream of the stop codon (**Figure 5**). This suggests that reverse transcription/integration of *KRTAP1-n* mRNA is unlikely to explain the pattern of *KRTAP1* concerted evolution, as the transcribed flanking regions of the gene would be expected to ‘hitch-hike’ with the coding regions through such a mechanism.

**Figure 5.**
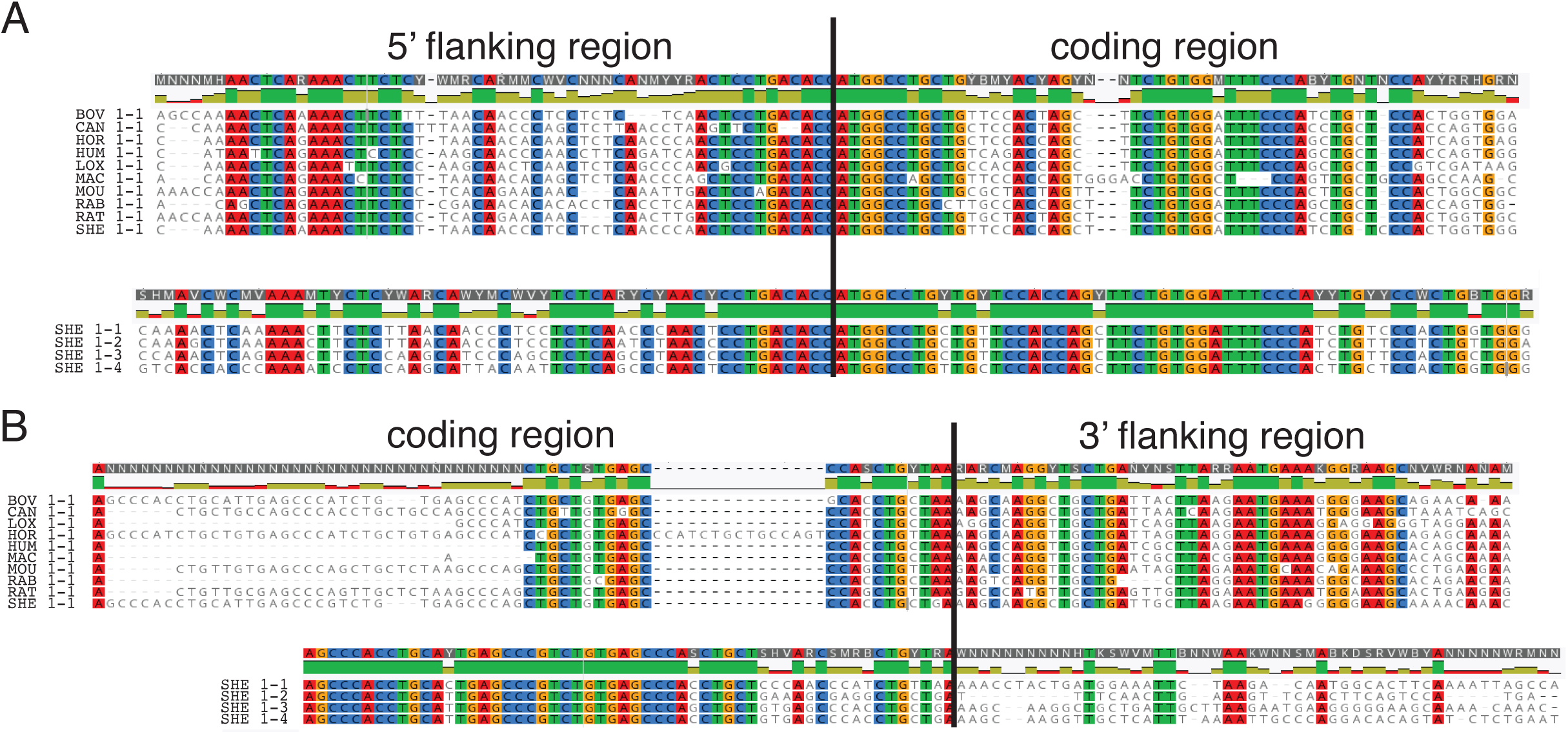
The switch between concerted and radiating evolution patterns is located close to the start/stop sites. **A**) Alignment of the region flanking the *KRTAP1-1* gene start site. The boundary between the 5’ flanking and coding regions is marked by a vertical line (followed by the ATG). Underneath is an alignment of the same region for all four *KRTAP1-n* sequences from sheep. Mismatches have a white background, conservation is indicated graphically above each alignment, and consensus sequences are shown at the top. **B)** As in **(A)**, except the region flanking the stop site is shown, with the vertical line marking the boundary between the coding and 3’ flanking regions (preceded by the stop codon).

We also considered whether the *KRTAP1-n* sequences might have arisen through a pure birth-and-death process by independent gene duplication events. However, we think this is improbable as it would require the same number of duplications to occur in at least seven of the species, and, independently, that each of these duplications would not involve any flanking sequence (including promoter and terminator sequences) and have inserted into the same site in each species.

### Gene conversion

Finally, we considered whether gene conversion could explain the pattern of *KRTAP1* repeat evolution. Gene conversion events within a genome that convert a section of one repeat to the sequence of another can create homogeneity (Chen, et al. 2007), and the degree of homogeneity depends on the relative rates of gene conversion and mutation (Teshima and Innan 2004; Harpak, et al. 2017). Our results imply that if gene conversion does occur, it is somehow restricted to the coding region. This pattern could occur if there is selective pressure to maintain a degree of intra-genome homogeneity between the repeat copies. If so, under the assumption that gene conversion occurs in both the coding and flanking regions, those events occurring in the flanking region will not have a selective advantage, while those occurring in the coding region will. Therefore, the probability of gene conversion events becoming fixed in the population will be greater for events that involve the coding region. There is considerable intra-genomic variation between *KRTAP1* repeats (**Figure 3**), but this incomplete level of homogenization can be explained by relatively infrequent gene conversion events and/or relative infrequent fixation of these events. Therefore, the sequence features of the *KRTAP1* repeats that we document here can all be accounted for by gene conversion coupled with selection.

### Evidence for gene conversion events in the *KRTAP1-n* repeats

Inspection of the *KRTAP1* coding region multiple sequence alignment provides evidence for tracts of gene conversion. Specifically, sites where there are mutations that are shared between copies within a species, but that differ between species, are frequently clustered together rather than scattered throughout the gene (**Figure 6**). Such patches of homogeneity are expected if there has been occasional, short-tract gene conversion events. The patches we observe are small, but are within the expected range for mammalian gene conversion events (Chen, et al. 2007). In addition, we collected population polymorphism data for *KRTAP1-n* sequences in sheep, as comprehensive sequence variation data are scarce in other species. For many of the sites that are polymorphic, the polymorphism is shared across some, or all, of the *KRTAP1-n* sequences (**Figure 7**). While we cannot rule out independent mutation events in each *KRTAP1* copy, we think that gene conversion is a more parsimonious explanation for this observation, particularly for the polymorphisms at synonymous sites. Gene conversion has also previously been suggested as an explanation for the pattern of polymorphism in the ovine *KRTAP1* genes (Rogers, et al. 1994). Collectively, our results suggest that the unusual evolutionary pattern of the *KRTAP1* repeats, where the coding region evolutionary dynamics are uncoupled from those of the flanking region, is the result of occasional short-tract gene conversion events that are selected for in the coding region but not the flanking regions, and that drive partial homogenization.

**Figure 6.**
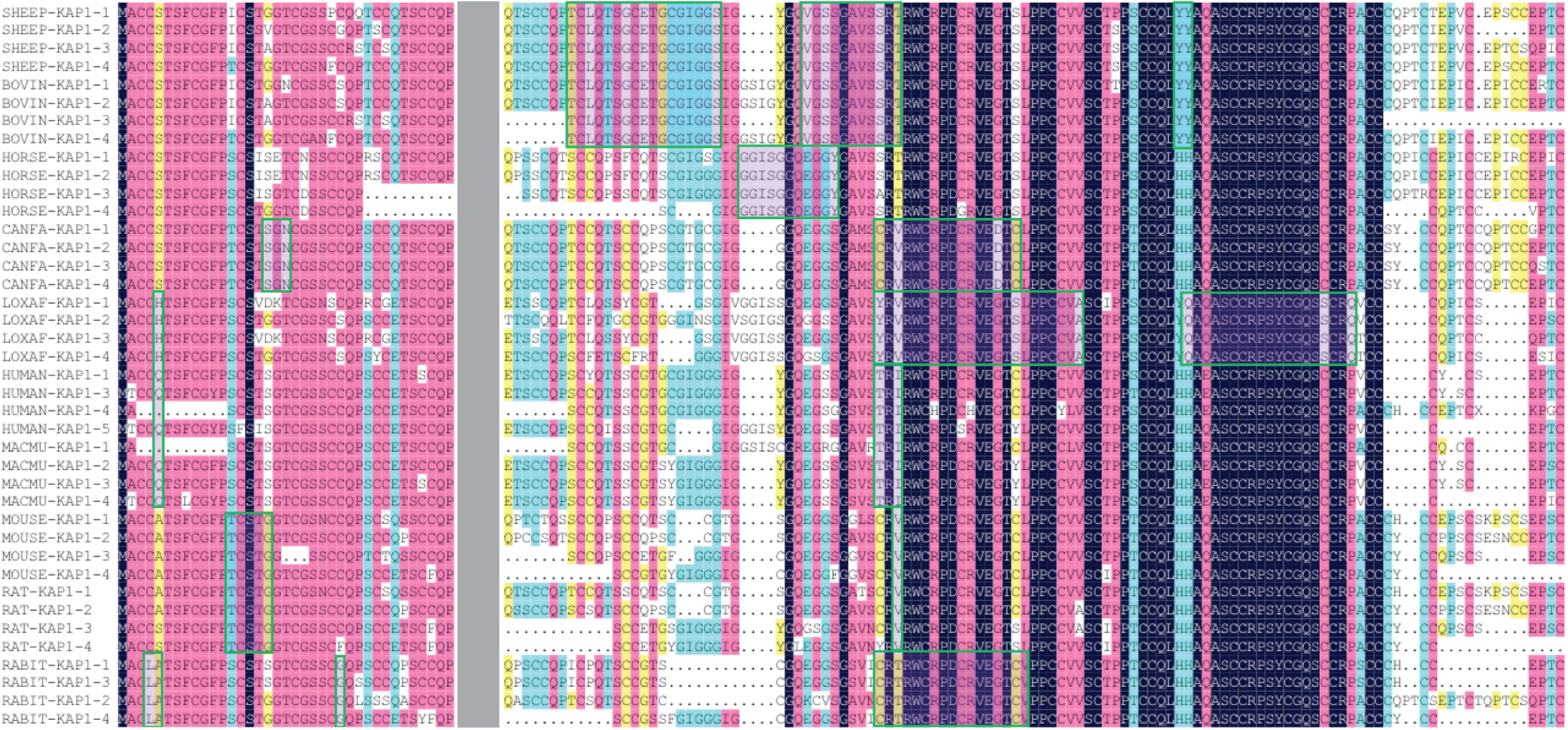
Evidence for short gene conversion tracts between *KRTAP1-n* sequences within species. Alignment of KAP1 amino acid sequences from the ten mammalian species. Amino acid tracts boxed in green represent sequences unique to a species or related species pairs. The grey vertical box represents the conserved decapeptide repeat sequences (which have been removed). Dots represent gaps in the alignment.

**Figure 7.**
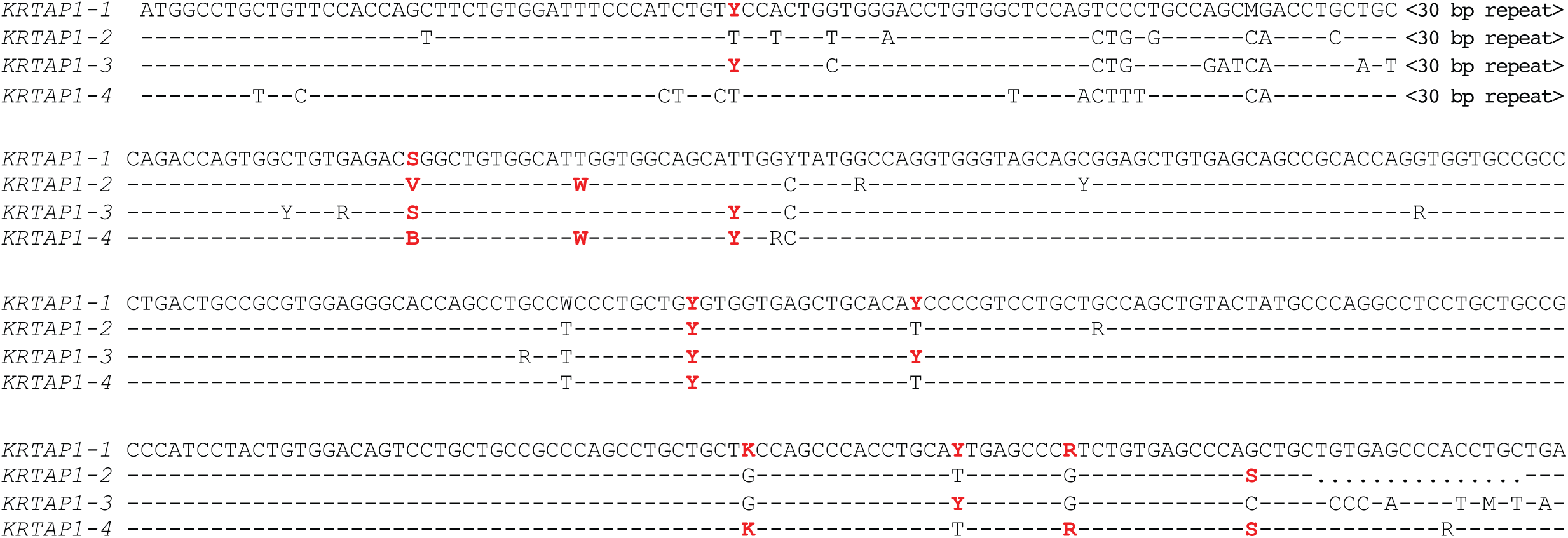
Shared polymorphisms between *KRTAP1-n* sequences in sheep. Alignment of the four sheep *KRTAP1-n* coding region sequences. Dashes represent nucleotides identical to the top sequence, and dots represent gaps. The 30 bp repeats are not shown, as the insertion/deletion positions cannot be precisely determined. Shared nucleotide substitutions between repeat copies are highlighted in red.

## Discussion

Here we have shown that *KRTAP1-n* genes are conserved as a block of four tandem repeats in mammalian species, and this suggests they derive from a relatively ancient gene-amplification event or events that probably pre-date mammalian speciation. The four tandem copies display a strong pattern of concerted evolution in the coding regions, yet the regions flanking show a normal radiating pattern of evolution. We suggest that this dichotomous pattern of evolution is not the result of purifying selection acting to retard changes to the amino acid sequence, but instead results from short gene conversion tracts that periodically homogenize sequences between the four *KRTAP1* genes within a genome.

The role of gene conversion is supported by two key pieces of evidence: 1) unique amino acid tracts that are shared by KAP1 copies within a species, but are unique to that species/group of related species; and 2) the possession of shared nucleotide variants between *KRTAP1* gene copies in sheep populations. These results extend previous reports of homogenization via ongoing short-tract gene conversion events in other protein coding genes (Noonan, et al. 2004; Lamping, et al. 2017).

We propose that gene conversion is being utilized as an unusual form of purifying selection that prevents accumulation of too much divergence between *KRTAP1* gene copies. We speculate that homogeneity of the *KRTAP1* coding sequences is beneficial as it enables the production of more homogenous components of the hair and wool fibre matrix, and thus potentially facilitates better associations with the keratin intermediate filaments. We cannot, however, rule out the possibility that individual *KRTAP1* repeats might have functional differences, the signal of which is overwhelmed by the concerted evolution signal from the majority of the gene. However, we note that, particularly in dogs, some of the *KRTAP1-n* genes are very similar in sequence. Therefore, we favour the explanation that *KRTAP1* concerted evolution results from ongoing, stochastic gene conversion events coupled with selection within the coding region against inter-repeat heterogeneity.

Purifying selection is evident in the *KRTAP1-n* coding regions, as the rate of synonymous change is about twice that of the non-synonymous rate (**Figure 4**). While this may seem to contradict the similarity in the synonymous and non-synonymous concerted evolution tree topologies, it can be simply explained by purifying selection acting on residues that are conserved between species, and thus not contributing to the synapomorphies that influence the tree topologies. Any gene conversion events that homogenize unfavourable amino acids will be selected against, thereby preventing deleterious mutations from spreading between copies. However, this same process also allows tolerable and advantageous amino acid changes to sweep through the copies (Dover 1982). The *KRTAP1-n* sequences from closely related species (i.e. human and macaque, rat and mouse, sheep and cattle) were not separated into different clades for most of the phylogenetic trees we generated (**Figures 3 and 4**). This suggests that the rate of homogenization is relatively slow, and insufficient to drive substantial homogeneity over the evolutionary time frames separating these species pairs. In this context, the shared polymorphisms that we observe in sheep (that are evidence for gene conversion events) are likely intermediate stages in the accumulation of homogenized *KRTAP1-n* sequences.

The sharp border between a concerted evolution pattern in the coding region and a radiating evolution pattern in the immediate flanking regions is striking. This can partially be explained by the selection for gene conversion events within the coding region, as we have proposed. However, it is intriguing to speculate that this may also be a consequence of differential expression between the *KRTAP1* genes that is mediated by copy-specific differences in the regulatory regions. Although not direct, some evidence for differential regulation of *KRTAP1-n* gene expression was found in two transcriptome studies looking for differentially expressed genes (Fan, et al. 2013; Chang, et al. 2014). If the *KRTAP1-n* genes do have functionally distinct roles, gene conversion events in the *KRTAP1* regulatory regions that perturb their differential regulation may be maladaptive and therefore selected against. Thus, selective pressure for coding region homogeneity versus regulatory region diversity, coupled with ongoing gene conversion, may be a powerful way to achieve the dichotomy in evolutionary patterns we observe. Clearly, a better understanding of the transcriptional regulation of the *KRTAP1* genes is required to address this hypothesis.

Gene conversion is frequently viewed through the lens of impeding sub-functionalization of gene duplicates. This view is consistent with the well characterized case of the opsin gene duplicates in primates, where there is a much stronger signal of gene conversion/concerted evolution in the introns, than in the exons (Shyue, et al. 1994; Hiwatashi, et al. 2011). The interpretation is that selection has largely rejected gene conversion events that include the coding (exon) regions, whilst allowing those occurring in the non-coding (intron) regions to spread in the population (Shyue, et al. 1994). This is the opposite of what we observe, and illustrates how gene conversion and selection can intersect to produce a constellation of evolutionary patterns: homogenization of the non-coding but not the coding regions in the opsin paralogs (Shyue, et al. 1994); homogenization of the coding but not the non-coding regions in the *KRTAP1* genes (this study); and homogenisation of both coding and non-coding regions equally in the ribosomal RNA gene repeats (Ganley and Kobayashi 2007).

The extent to which gene conversion acts to homogenize gene duplicates remains controversial (Gao and Innan 2004; Casola, et al. 2012; Harpak, et al. 2017). Furthermore, even in examples where recurrent gene conversion events can be detected, they are often not sufficient to produce a strong concerted evolution pattern (Petronella and Drouin 2011, 2014). There are two potential explanations for why such a strong pattern of concerted evolution is observed in the case the *KRTAP1* genes, despite the relatively high levels of divergence between copies. First, unlike many of the examples that have aroused controversy (Gao and Innan 2004; Casola, et al. 2012; Harpak, et al. 2017), the *KRTAP1-n* repeats are tandemly-arranged. Proximity effects as a consequence of tandem arrangement may increase the chances of unequal alignment of the repeats during DNA repair-based homologous recombination compared to dispersed repeats, and thus may increase the chances of inter-repeat gene conversion events. However, this does not explain examples where tandemly repeated paralogs do not show a strong concerted evolution pattern (Nei, et al. 2000; Perina, et al. 2011). A second explanation relates to the imperfect decapeptide tandem repeat motif found in the coding region. Variation in the copy number of decapeptide repeats between *KRTAP1* genes is possibly the result of unequal recombination (Liao and Weiner 1995; Ganley and Scott 1998; Morrill, et al. 2016). If so, the *KRTAP1* genes may harbour a recombination hotspot that drives both decapeptide repeat copy number variation and gene conversion at higher than average levels.

Repeats are ubiquitous denizens of eukaryote genomes, where they exist in different forms (coding, non-coding) and organizations (tandem, dispersed). Our results add to the growing list of examples that illustrate how different molecular and evolutionary processes can impinge on repeats to structure their sequences and create distinctive patterns of evolution (Shyue, et al. 1994; Noonan, et al. 2004; Ganley and Kobayashi 2007; Storz, et al. 2007; Hiwatashi, et al. 2011; Lamping, et al. 2017). However, it is unclear how widespread these sorts of evolutionary dynamics are for eukaryotic gene repeats, largely because the patterns of evolution have not been investigated for the vast majority of multi-gene families. The increasing availability of high quality genome sequences for a wide range of eukaryotes puts us in an excellent position to determine, on a much more systematic and wide-ranging basis, the patterns of repeat sequence dynamics and evolution. This will, in turn, make it clear whether the impact of recombination on the *KRTAP1*s is unusual, or highlights a common mechanism to finely scale patterns of homogeneity and divergence between repeat copies over time.

## Acknowledgements

This work was supported by a Marsden Fund award (14-MAU-053) to ARDG, an AGMARDT Postdoctoral Fellowship to HG, and a Vernon Willey Trust Fellowship to HZ.

