## Supplementary Materials for "Contrasting patterns of coding and flanking region evolution in mammalian keratin associated protein-1 genes"

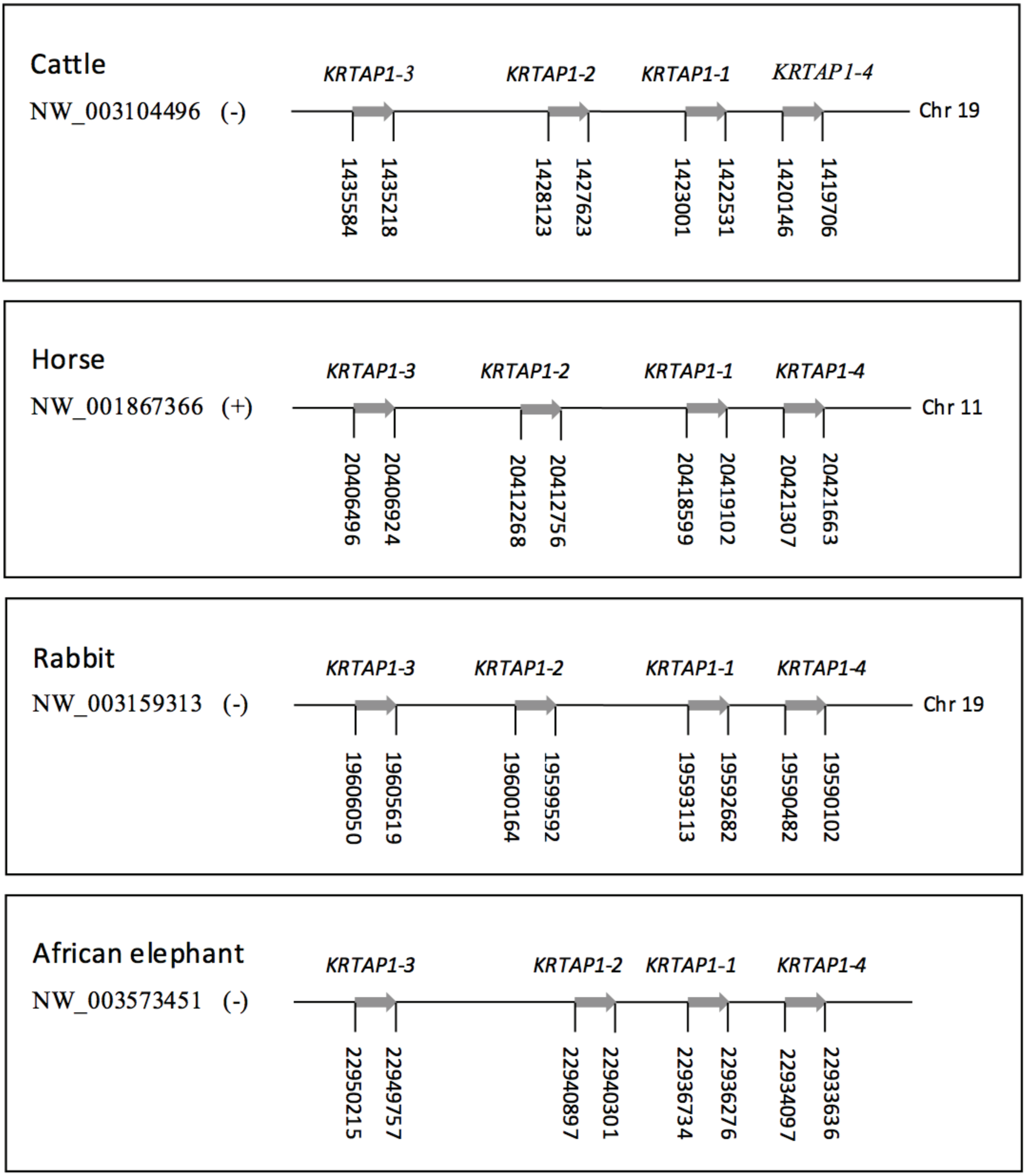


**Figure S1. Newly reported *KRTAP1* organization for four mammalian species.**
The genomic organization of the *KRTAP1-n* repeats is shown for four mammalian species that it has not been previously reported for. Genbank accession numbers are indicated to the left, the strand is indicated in parentheses, and the start and stop positions are indicated below the genes. The chromosome is indicated to the right where known.
