## Supplementary Materials for "Contrasting patterns of coding and flanking region evolution in mammalian keratin associated protein-1 genes"

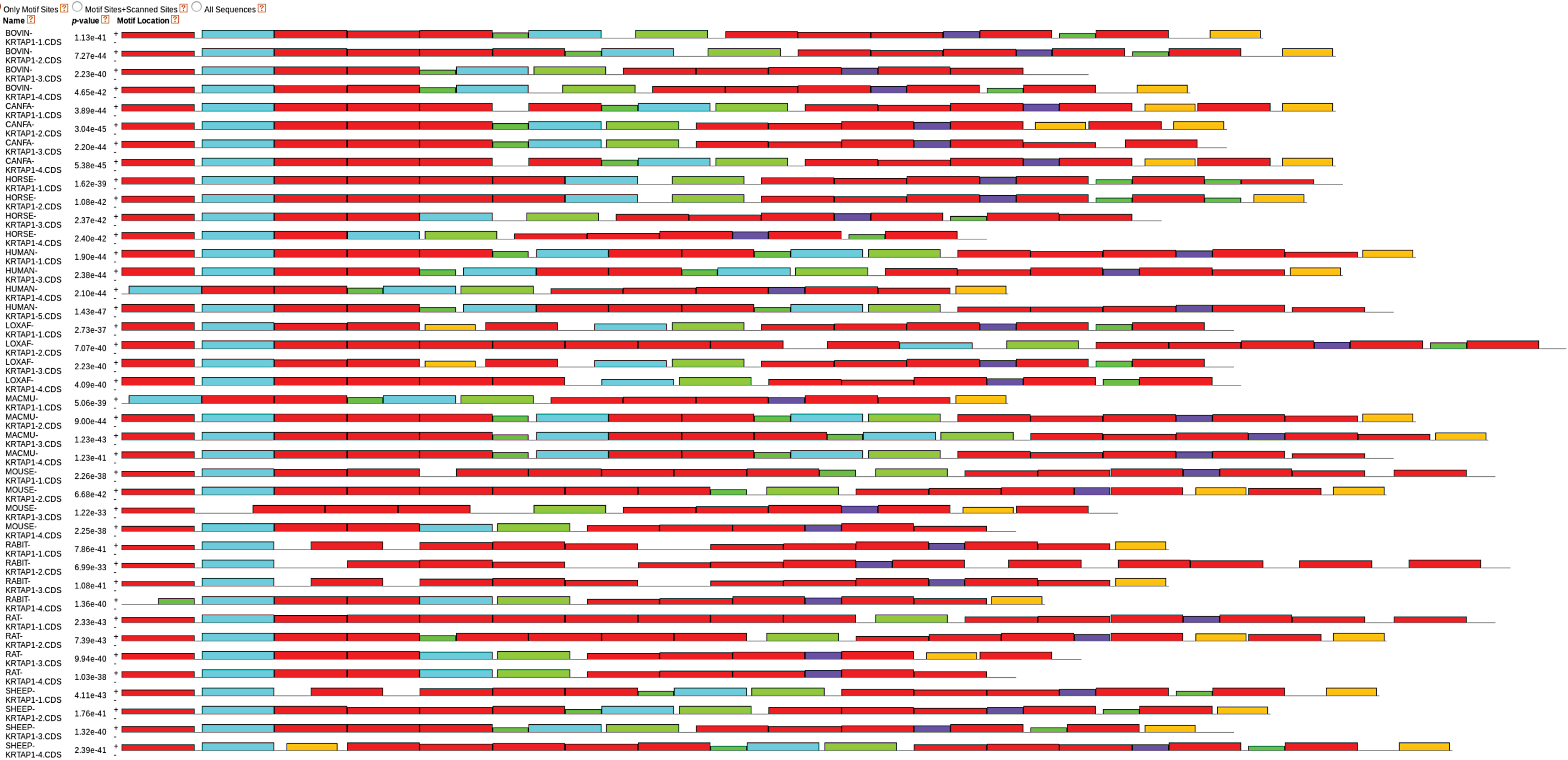


**Figure S2. MEME identifies copies of the 30 bp intragenic *KRTAP1* repeat at both ends of the gene.**
MEME output showing the positions of 30 bp repeat motifs amongst all *KRTAP1-n* copies from all ten mammalian species. The red boxes represent the previously identified 30 bp repeat that is present at the 5' and 3' ends of the gene and varies in copy number between repeat copies within and between species. Other colored boxes are other motifs detected by MEME, and are predominantly motifs shared between rather than within genes.
