## Supplementary Materials for "Contrasting patterns of coding and flanking region evolution in mammalian keratin associated protein-1 genes"

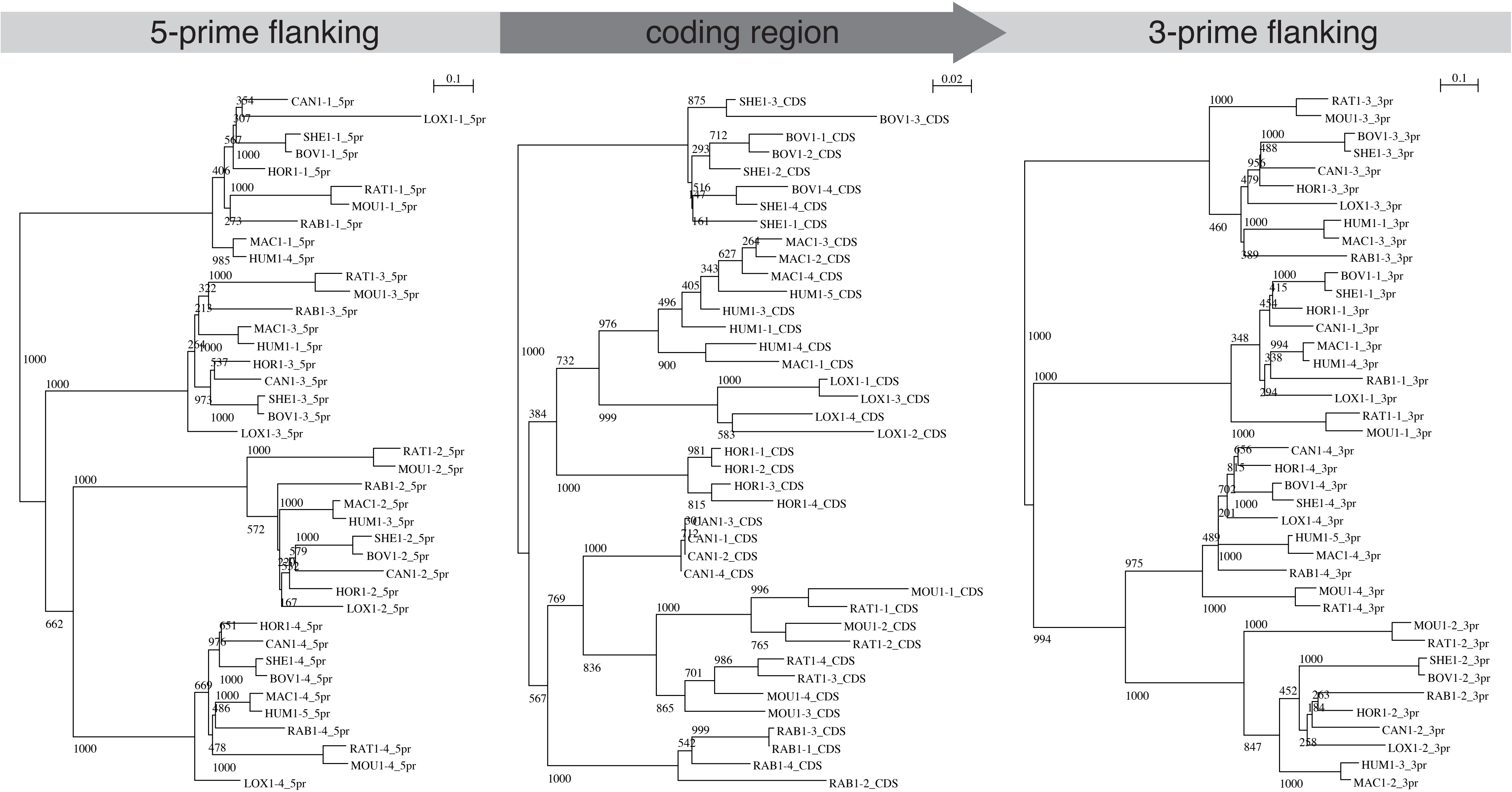


**Figure S3. Phylogenies from more relaxed alignment criteria still show a *KRTAP1* coding region-specific concerted evolution pattern.** Phylogenetic trees were generated for mammalian *KRTAP1-n* 5'-flanking, coding, and 3'-flanking regions using PhyML with multiple sequences alignments that included more poorly aligning regions than **Figure 3**. Species are indicated by three-letter abbreviations followed by "1" for *KRTAP1*, number corresponding to *KRTAP1-n* repeat number, and the region the sequence is from. The 5' and 3' flanking phylogenies group by repeat number, whereas the coding region phylogeny tends to group by species. The numbers on the nodes indicate bootstrap support, and substitution rates are indicated above.
