## Supplementary Materials for "Contrasting patterns of coding and flanking region evolution in mammalian keratin associated protein-1 genes"

**Table S1.** *KRTAP1* sequences from the genomes of the ten mammalian species used in this study.

| Species | Gene | GenBank Accession number | Open reading frame | Strand |
|---|---|---|---|---|
| Sheep | *KRTAP1-1* | NC_019468.2 | 40766938 – 40767417 | – |
| | *KRTAP1-2* | NC_019468.2 | 40771911 – 40772378 | – |
| | *KRTAP1-3* | NC_019468.2 | 40779170 – 40779617[a] | – |
| | *KRTAP1-4* | NC_019468.2 | 40764049 – 40764578 | – |
| Human | *KRTAP1-4* | NT_010783.15 | 4460117 – 4460482 | – |
| | *KRTAP1-3* | NT_010783.15 | 4464722 – 4465225 | – |
| | *KRTAP1-1* | NT_010783.15 | 4471268 – 4471801 | – |
| | *KRTAP1-5* | NT_010783.15 | 4457035 – 4457559 | – |
| Macaque | *KRTAP1-1* | NC_027908.1 | 46855931 – 46856296 | – |
| | *KRTAP1-2* | NC_027908.1 | 46862830 – 46863363 | – |
| | *KRTAP1-3* | NC_027908.1 | 46870332 – 46870895 | – |
| | *KRTAP1-4* | NC_027908.1 | 46852802 – 46853326 | – |
| Dog | *KRTAP1-1* | NW_003726075.1 | 3461261 – 3461761 | + |
| | *KRTAP1-2* | NW_003726075.1 | 3453667 – 3454122 | + |
| | *KRTAP1-3* | NW_003726075.1 | 3447783 – 3448238 | + |
| | *KRTAP1-4* | NW_003726075.1 | 3464126 – 3464626 | + |
| Mouse | *KRTAP1-1* | NT_165773.2 | 10985451 – 10986017 | – |
| | *KRTAP1-2* | NT_165773.2 | 10993187 – 10993708 | – |
| | *KRTAP1-3* | NT_165773.2 | 11000033 – 11000443 | – |
| | *KRTAP1-4* | NT_165773.2 | 10982986 – 10983354 | – |
| Rat | *KRTAP1-1* | NW_047339.1 | 1328764 – 1329330 | – |
| | *KRTAP1-2* | NW_047339.1 | 1335870 – 1336391 | – |
| | *KRTAP1-3* | NW_047339.1 | 1342514 – 1342909 | – |
| | *KRTAP1-4* | NW_047339.1 | 1326287 – 1326655 | – |
| Cattle | *KRTAP1-1* | NW_003104496.1 | 1422531 – 1423001 | – |
| | *KRTAP1-2* | NW_003104496.1 | 1427623 – 1428123 | – |
| | *KRTAP1-3* | NW_003104496.1 | 1435218 – 1435584 [b] | – |
| | *KRTAP1-4* | NW_003104496.1 | 1419706 – 1420146 | – |
| Horse | *KRTAP1-1* | NW_001867366.1 | 20418599 – 20419102 | + |
| | *KRTAP1-2* | NW_001867366.1 | 20412268 – 20412756 | + |
| | *KRTAP1-3* | NW_001867366.1 | 20406496 – 20406924 | + |
| | *KRTAP1-4* | NW_001867366.1 | 20421307 – 20421663 | + |
| Elephant | *KRTAP1-1* | NW_003573451.1 | 22936276 – 22936734 | – |
| | *KRTAP1-2* | NW_003573451.1 | 22940301 – 22940897 | – |
| | *KRTAP1-3* | NW_003573451.1 | 22949757 – 22950215 | – |
| | *KRTAP1-4* | NW_003573451.1 | 22933636 – 22934097 | – |
| Rabbit | *KRTAP1-1* | NW_003159313.1 | 19592682 – 19593113 | – |
| | *KRTAP1-2* | NW_003159313.1 | 19599592 – 19600164 | – |
| | *KRTAP1-3* | NW_003159313.1 | 19605619 – 19606050 | – |
| | *KRTAP1-4* | NW_003159313.1 | 19590102 – 19590482 | – |

[a] the sheep genome assembly accession is missing 11 nucleotides. We used Genbank accession X02925.1 (which has a complete coding region) for the coding sequence, and the genome assembly for the flanking region sequences.

[b] The end of the coding region is missing from the cattle genome assembly. We used Genbank accession XM_001255391.4 (which has a putatively complete coding region) for the coding sequence, and the genome assembly for the flanking region sequences.
