## Supplementary Materials for "Contrasting patterns of coding and flanking region evolution in mammalian keratin associated protein-1 genes"

**Table S2.** No evidence for codon bias explaining the *KRTAP1* coding region pattern of concerted evolution

| Human variant | Mouse variant | Human codon usage[a] | | Mouse codon usage[a] | | Support[b] |
|---|---|---|---|---|---|---|
| GGC | GGT | 22.3 | 10.6 | 21.2 | 11.4 | |
| AGG | AGA | 11.9 | 12 | 12.2 | 12.1 | |
| CCA | CCT | 17.3 | 17.9 | 17.3 | 18.4 | |
| CGT | CGC | 4.5 | 10.5 | 4.7 | 9.4 | |
| GG(T/C)[c] | GGC | 10.6 | 22.3 | 11.4 | 21.2 | |
| TG(C/T)[c] | TGT | 12.2 | 10.6 | 12.3 | 11.4 | |
| CAC | CAT | 15.2 | 11.1 | 15.3 | 10.6 | |
| CGC | CGT | 10.5 | 4.5 | 9.4 | 4.7 | |
| CC(G/A)[c] | CCA | 7.3 | 17.3 | 6.2 | 17.3 | |

a. Value on the left is the codon usage value for the human variant codon; that on the right for the mouse variant codon

b. Dark colors = both codons support/neutral for CAI explaining (green)/not explaining (red) concerted evolution pattern; light colors is where support is opposite for each codon with color representing the dominant effect (in terms of CAI magnitudes)

c. Difference is present in two of the four *KRTAP1* copies. Codon usage values on the left represent the human variant that differs from the mouse
